# Exact solutions of linear reaction-diffusion processes on a uniformly growing domain: Criteria for successful colonization

**DOI:** 10.1101/011122

**Authors:** Matthew J Simpson

**Affiliations:** Mathematical Sciences, Queensland University of Technology, Brisbane, Australia.

## Abstract

Many processes during embryonic development involve transport and reaction of molecules, or transport and proliferation of cells, within growing tissues. Mathematical models of such processes usually take the form of a reaction-diffusion partial differential equation (PDE) on a growing domain. Previous analyses of such models have mainly involved solving the PDEs numerically. Here, we present a framework for calculating the exact solution of a linear reaction-diffusion PDE on a growing domain. We derive an exact solution for a general class of one-dimensional linear reaction-diffusion process on 0 < *x* < *L*(*t*), where *L*(*t*) is the length of the growing domain. Comparing our exact solutions with numerical approximations confirms the veracity of the method. Furthermore, our examples illustrate a delicate interplay between: (i) the rate at which the domain elongates, (ii) the diffusivity associated with the spreading density profile, (iii) the reaction rate, and (iv) the initial condition. Altering the balance between these four features leads to different outcomes in terms of whether an initial profile, located near *x* = 0, eventually overcomes the domain growth and colonizes the entire length of the domain by reaching the boundary where *x* = *L*(*t*).

## Introduction

Developmental processes are often associated with transport and reaction of molecules, or transport and proliferation of cells, within growing tissues [1, 2]. For example, the development of biological patterns, such as animal coat markings, is thought to arise due to the coupling between an activator-inhibitor Turing mechanism and additional transport induced by tissue growth [3–7]. Within the mathematical biology literature, there is an increasing awareness of the importance of incorporating domain growth into mathematical models of various biological processes including morphogen gradient formation [8] and models of collective cell spreading [9]. In addition to considering particular biological applications, other studies have focused on examining more theoretical questions associated with reactive transport processes on growing domains. Most notably, several previous studies have examined the relationship between discrete random walk models and associated continuum partial differential equation (PDE) descriptions [10–14].

One particular biological application where transport and reaction (proliferation) of cells takes place on a growing domain is the development of the enteric nervous system (ENS) [15–21]. This developmental process involves neural crest precursor cells entering the oral end of the developing gut. Individual precursor cells migrate and proliferate, which results in the formation of a moving front of precursor cells which travels towards the anal end of the developing gut. This colonization process is complicated by the fact that the gut tissues elongate simultaneously as the cell front moves [17]. Normal development requires that the moving front of precursor cells reaches the anal end of the developing tissue. Abnormal development is thought to be associated with situations where the moving front of cells fails to completely colonize the growing gut tissue [17].

One of the first mathematical models of ENS development, described by Landman et al. [22], is a PDE description of the migration and proliferation of a population of precursor cells on a uniformly growing tissue. In particular, Landman et al. [22] use their model to mimic ENS development by considering an initial condition where the population of precursor cells is initially confined towards one end of the domain. Landman et al. [22] solve the governing PDE numerically and use these numerical solutions to explore whether the population of cells can colonize the entire length of the growing domain within a certain period of time. In particular, Landman et al. [22] highlights an important interaction between: (i) the initial distribution of cells; (ii) the migration rate of cells; (iii) the proliferation rate of cells; and (iv) the growth rate of the underlying tissue. Landman et al. [22] explore the relationship between these four factors using an approximate numerical solution of the PDE model. These previous numerical results suggest that successful colonization requires: (i) that the initial length of colonization must be sufficiently large, (ii) that the migration rate of cells is sufficiently large, (iii) that the proliferation rate of cells is sufficiently large, and (iv) that the growth rate of the underlying tissue is sufficiently small.

In addition to presenting numerical solutions, Landman et al. [22] also presents analysis for the special case where there is no cell diffusion. This analysis involves solving a simplified hyperbolic PDE model using the method of characteristics. While this analysis offers useful insight, Landman et al. [22] does not provide any exact solutions for the case where diffusive transport is included.

The focus of the present work is to consider a linear reaction-diffusion process on a growing domain with a view to obtaining an exact solution of the associated PDE. After transforming the PDE to a fixed domain we obtain a PDE with variable coefficients. The variable coefficient PDE is simplified using an appropriate transformation which enables us to obtain an exact solution using separation of variables. While our strategy for obtaining an exact solution is quite general, we present specific results for linear and exponentially elongating domains. After verifying the accuracy of our exact solutions using numerical approximations, we summarise our results in terms of a concise condition that can be used to distinguish between successful or unsuccessful colonization. We conclude this study by acknowledging the limitations of our analysis, and we outline some further extensions of our approach which could be implemented in future studies.

## 1 Materials and methods

### 1.1 Mathematical model

We consider a linear reaction-diffusion process on a one-dimensional domain, 0 < *x* < *L*(*t*), where *L*(*t*) is the increasing length of the domain. Domain growth is associated with a velocity field which causes a point at location *x* to move to *x* + *v*(*x*,*t*)*τ* during a small time period of duration *τ* [22]. By considering the expansion of an element of initial width Δ*x*, we can derive an expression relating *L*(*t*) and *v*(*x*,*t*), which can be written as

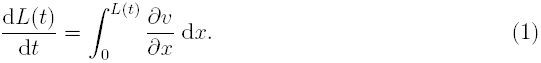

Like others [3, 4, 22], we consider uniform growth conditions where 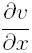 is independent of position, but potentially depends on time, *t*, so that we have 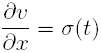. Combining this definition with Equation (1) gives:

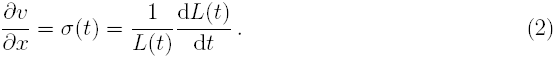

Without loss of generality, we assume that the domain elongates in the positive *x*-direction with the origin fixed, so that *v*(0,*t*) = 0. Integrating Equation (2) gives

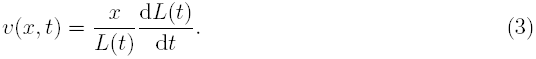

We now consider conservation of mass of some density function, *C*(*x*, *t*), assuming that the population density function evolves according to a linear reaction-diffusion mechanism. The associated conservation statement on the growing domain can be written as

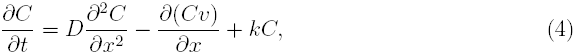

on 0 < *x* < *L*(*t*), where *D* > 0 is the diffusivity, *k* is the production rate and *v* is the velocity associated with the underlying domain growth, given by Equation (3). We note that setting *k* > 0 represents a source term which is relevant to ENS development since the precursor cells proliferate [16–18,20]; however, our approach can also be used to study decay processes by setting *k* < 0.

To solve Equation (4) we must specify initial conditions and boundary conditions. Motivated by Landman et al. [22], we choose

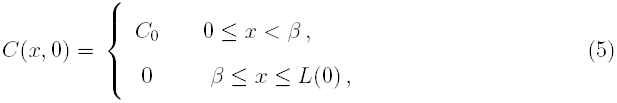

which corresponds to some initial length of the domain, 0 ≤ *x* < *β*, being uniformly colonized at density *C*_0_, with the remaining portion of the domain being uncolonized. We suppose that we have zero diffusive flux conditions at both boundaries, 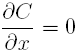 at *x* = 0 and *x* = *L*(*t*), and we now seek to find an exact solution, *C*(*x*,*t*).

## 2 Results

### 2.1 Exact solution

The first step in our solution strategy is to transform the spatial variable to a fixed domain, 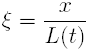 [3, 4, 10–12, 22], giving

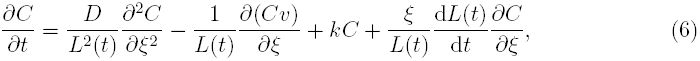

on 0 < *ξ* < 1. Recalling that 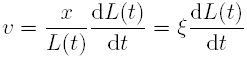, we re-write Equation (6) as

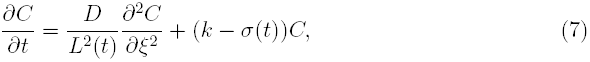

where, in the transformed coordinates, the impact of domain growth manifests in two different ways:

1. the coefficient of the diffusive transport term is inversely proportional to *L*^2^(*t*), and hence decreases with time, and
2. the addition of a source term, *−Cσ*(*t*), represents dilution associated with the expanding domain.

Following Crank [23] we re-scale time,

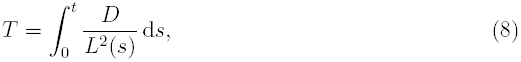

giving

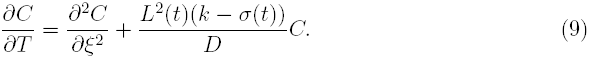

Equation (8) gives a relationship between the original time variable, *t,* and the transformed variable, *T*, which means that we can write Equation (9) as

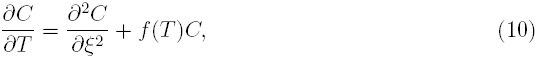

whose solution, with zero diffusive flux conditions at both boundaries, can be obtained by applying separation of variables [23], giving

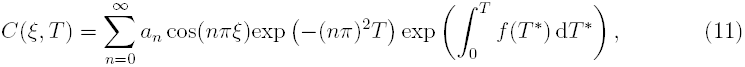

where *n* ∈ ℕ^0^. Our exact solution for *C*(*ξ*,*T*) can be re-written in terms of the original coordinates, giving *C*(*x*,*t*). The Fourier coefficients, *a*_*n*_, can be chosen to ensure that the exact solution satisfies the initial condition, given by Equation (5). Our framework for finding *C*(*x*,*t*) is quite general and does not depend on any particular form of the initial condition. We now present the details for a few relevant choices of *L*(*t*).

#### 2.1.1 Case 0: Non-growing domain

Before we present results for a growing domain it is instructive to consider the solution of Equation (4), with the same initial condition and boundary conditions, on a non-growing domain, 0 < *x* < *L*. With *L*(*t*) = *L*, we have *σ*(*t*) = 0 and 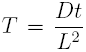. Later, when we compare the solution of Equation (4) on a growing domain with the solution on a non-growing domain, it will be useful to recall that on a non-growing domain, as *t* → ∞, we have *T* → ∞, since *D* > 0 and *L* > 0. On the non-growing domain the solution of Equation (4) can be written as

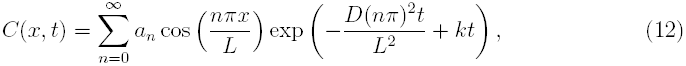

where 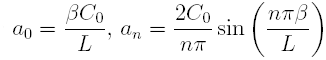 and *n* ∈ ℕ^+^.

#### 2.1.2 Case 1: Exponential domain growth

With *L*(*t*) = *L*(0)exp (*αt*), we have *σ*(*t*) = *α* and 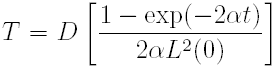, for which we note that as *t* → ∞, we have 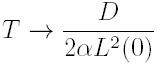, since *α* > 0. This limiting behavior is different to the limiting behavior under non-growing conditions. For an exponentially-elongating domain, the solution of Equation (4) can be written as

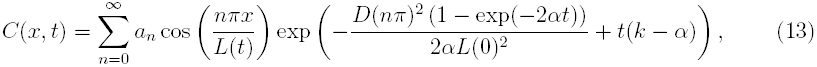

where 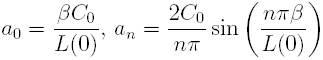 and *n* ∈ ℕ^+^.

#### 2.1.3 Case 2: Linear domain growth

With *L*(*t*) = *L*(0) + *bt*, we have 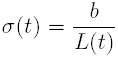 and 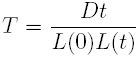 for which as *t* → ∞, we have 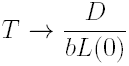, since *D* > 0 and *b* > 0. For a linearly-elongating domain, the solution of Equation (4) can be written as

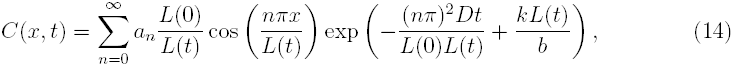

where 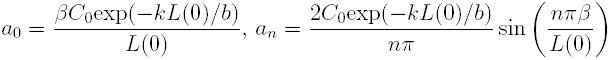 and *n* ∈ ℕ^+^.

### 2.2 Comparison of exact and numerical solutions

We now present some examples to highlight key features of the model. First we compare plots of *C*(*x*,*t*) generated using the exact solution with plots of *C*(*x*,*t*) computed numerically.To generate the numerical approximations we discretise Equation (7) using a central finite difference approximation on a uniformly discretized domain, 0 < *ξ* < 1, with uniform mesh spacing *δξ*. The resulting system of coupled ordinary differential equations is integrated through time using a backward Euler approximation with uniform time steps of duration *δt*. At each time step the resulting system of tridiagonal linear equations is solved using the Thomas algorithm [24]. All numerical results presented correspond to choices of *δξ* and *δt* such that the numerical results are grid-independent.

**Figure 1:**
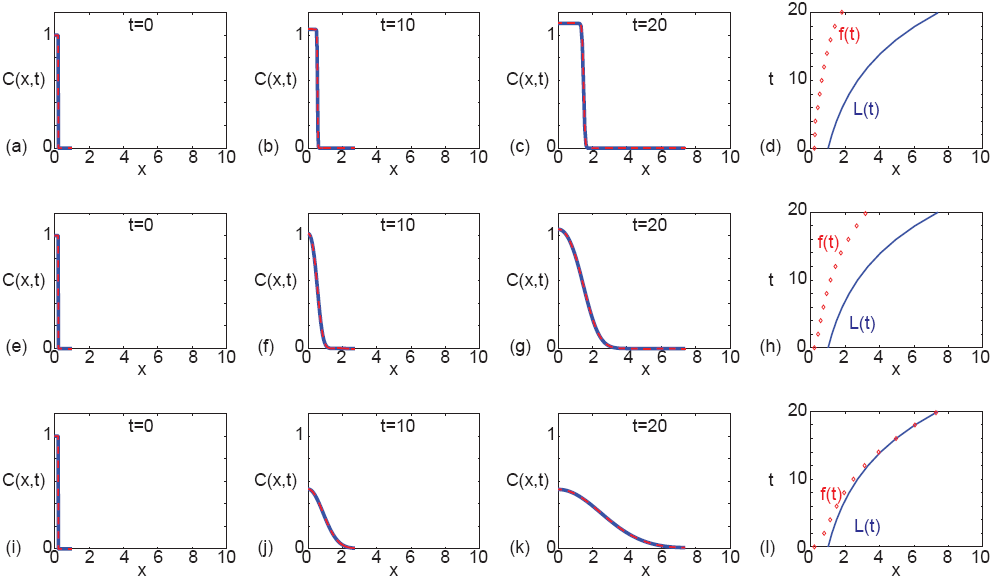
Comparison of exact and numerical solutions, exploring the influence of varying the diffusivity, *D*. All results correspond to an exponentially-elongating domain, *L*(*t*) = *L*(0)exp(*αt*), with *L*(0) = 1 and *α* = 0.1. The initial condition is given by Equation (5) with *β* = 0.2 and *C*_0_ = 1. In all cases we consider a linear source term with *k* = 0.105. Results in (a)–(d) correspond to *D* = 0.00001, results in (e)–(h) correspond to *D* = 0.001, and results in (i)–(l) correspond to *D* = 0.01. For all three sets of parameter combinations we show the solution at *t* = 0,10 and *t* = 20, as indicated. The exact solutions, presented in (a)–(c), (e)–(g) and (i)–(k) (solid blue), correspond to Equation (13), where we truncate the infinite sum after 1000 terms. The numerical solutions, presented in (a)–(c), (e)–(g) and (i)–(k) (dashed red), are numerical approximations of Equation (7) with *δξ* = 0.001 and *δt* = 0.001. The space-time diagrams summarising the time evolution of the length of the domain, *L*(*t*), and the position of the front of the *C*(*x*,*t*) density profile, *f*(*t*), given in (d), (h) and (l), are constructed by defining *f*(*t*) to be the position where *C*(*x*,*t*) = 0.01.

Results in Fig. 1A-C compare exact and numerical solutions on an exponentially-growing domain at *t* = 0, 10 and 20, and we see that the exact and numerical solutions are indistinguishable. A summary of the properties of the solutions in the interval 0 ≤ *t* ≤ 20 is given in a space-time diagram in Fig. 1D, which compares the length of the domain, *L*(*t*), and the position of the front, *f*(*t*). Here, we define the position of the front to be the spatial location where *C*(*x*,*t*) = 0.01. This means that we have *f*(0) = *β*. Comparing *L*(*t*) and *f*(*t*) in Fig. 1D indicates that the *C*(*x*,*t*) profile moves in the positive *x*-direction as time increases; however, the distance between *L*(*t*) and *f*(*t*) increases with time such that the *C*(*x*,*t*) profile does not colonize the domain by *t* = 20.

Results in Fig. 1E-G correspond to the same initial condition and parameters used in Fig. 1A-C except that we increased the diffusivity, *D*. Comparing results in Fig. 1E-G with the solutions in Fig. 1A-C indicates that the front moves faster with an increase in *D*, as we might anticipate. However, the summary of the time evolution of *L*(*t*) and *f*(*t*) in Fig. 1H confirms that the increase in *D* is insufficient for colonization to occur by *t* = 20. In contrast, the results in Fig. 1I-K correspond to the same initial condition and parameters as in Fig. 1E-G except that we have further increased *D*. This time we see that the front reaches *L*(*t*), and we have full colonization after *t* ≈ 16.

**Figure 2:**
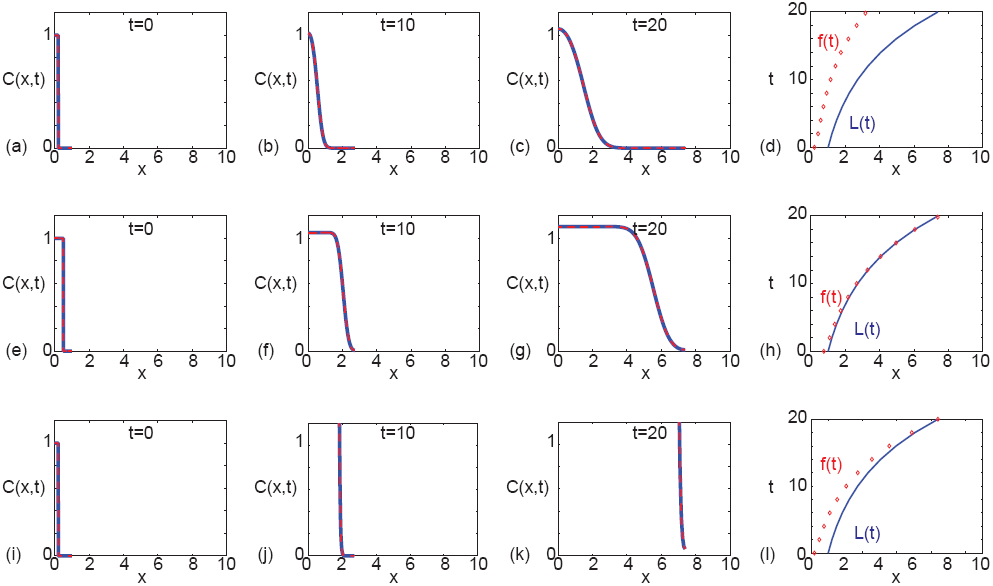
Comparison of exact solutions and numerical approximations for different values of β and *k*. All results correspond to an exponentially-elongating domain, *L*(*t*) = *L*(0)exp(*αt*), with *L*(0) = 1 and α = 0.1. The initial condition is given by Equation (5) with *C*_0_ = 1, and in all cases we set *D* = 1×10^-3^. Results in (a)–(d) correspond to a narrow initial condition, α = 0.2, with *k* = 0.105. Results in (e)–(h) correspond to a wide initial condition, β = 0.75, with *k* = 0.105. Results in (i)–(l) correspond to a narrow initial condition, β = 0.2, with *k* = 1.705. For each set of parameter combinations we show the solution at *t* = 0, 10 and *t* = 20, as indicated. The exact solutions, presented in (a)–(c), (e)–(g) and (i)–(k) (solid blue), correspond to Equation (13), where we truncate the infinite sum after 1000 terms. The numerical solutions, presented in (a)–(c), (e)–(g) and (i)–(k) (dashed red), correspond to are numerical approximations of Equation (7) with δξ = 0.001 and *δt* = 0.001. The space–time diagrams summarising the time evolution of the length of the domain, *L*(*t*), and the position of the front of the *C*(*x, t*) density profile, *f*(*t*), given in (d), (h) and (l), are constructed by defining *f*(*t*) to be the position where *C*(*x, t*) = 0.01.

To further explore the competition between various processes in the model we compare some additional exact and numerical solutions in Fig. 2, where again we see that in all cases considered, the numerical solutions are visually indistinguishable from the exact solutions. The set of results in Fig. 2A-D is identical to the set of results shown previously in Fig. 1E-H, which corresponds to a case where the domain does not become fully colonized within the interval 0 *≤ t* ≤ 20. We present a second set of results, in Fig. 2E-H, which are identical to those in Fig. 2A-D except for a change in the initial condition. We note that the initial condition in Fig. 2A-D corresponds to *C*(*x*, 0) = 1 for 0 ≤ *x* < 0.2 and *C*(*x*, 0) = 0 for 0.2 ≤ *x* ≤ 1, whereas the initial condition in Fig. 2E-H corresponds to *C*(*x*, 0) = 1 for 0 ≤ *x* < 0.75 and *C*(*x*, 0) = 0 for 0.75 ≤ *x* ≤ 1. The situation in Fig. 2A-D leads to unsuccessful colonization by *t* = 20 whereas the situation in Fig. 2E-H leads to successful colonization after *t* ≈ 14. A third set of results, in Fig. 2I-L, are identical to those in Fig. 2A-D except for a change in the production term *k*. For *k* = 0:105, profiles in Fig. 2A-D do not colonize the growing domain by *t* = 20. In contrast, when we increase the production to *k* = 1.705, profiles in Fig. 2I-L indicate that colonization occurs after *t* ≈ 20.

Although all results presented in Fig. 1 and Fig. 2 correspond to an exponentially-growing domain, we also generated exact and numerical results for a linearly elongating domain (not shown), and we note two key outcomes. First, similar to the results in Fig. 1 and Fig. 2, we found that the exact solution and the numerical solutions compare very well. Second, we found that altering the initial condition, *D*, *k* and the growth rate, *b*, could affect whether or not the system colonized within a specified time interval.

### 2.3 Criteria for colonization

Now that we have derived and validated exact solutions describing a linear reaction-diffusion process on a growing domain we can use the new solution to write down a condition which can be used to distinguish between situations which lead to successful colonization from situations which lead to unsuccessful colonization. For our initial condition, given by Equation (5), we aim to identify whether the spreading density profile, *C*(*x*,*t*), ever reaches the boundary, *x* = *L*(*t*), by some threshold time *t*^⋆^. To explore this we must examine the quantity *C*(*L*(*t*^⋆^),*t*^⋆^) by substituting *x* = *L*(*t*^⋆^) and *t* = *t*^⋆^ into Equation (11). Having evaluated this quantity, we test whether *C*(*L*(*t*^⋆^),*t*^⋆^) > *ε*, in which case we have successful colonization by time *t*^⋆^. Alternatively, if *C*(*L*(*t*^⋆^), *t*^⋆^) < *ε*, we have unsuccessful colonization by time *t*^⋆^. Here *ε* is some user-defined small tolerance. For example, to interpret the results in Fig. 1 and Fig. 2, we set *ε* = 0.01 to determine the position of the front, and this choice of *ε* could be used to make a distinction between successful and unsuccessful colonization in other applications.

We now demonstrate how our results are sensitive to the choice of ɛ. If we choose a slightly larger tolerance, say *ɛ* = 0:015, our conclusions about the results in Fig. 1 are slightly different. With *ɛ* = 0.015, our conclusion about the situations in Fig. 1A-D and Fig. 1E-H remains unchanged and colonization never occurs. However, for the parameter combination in Fig. 1I-L, the position of the moving front, according to the larger tolerance, takes a longer period of time to reach *x* = *L*(*t*). Instead of reaching *x* = *L*(*t*) by *t* ≈ 16 with *ɛ* = 0.01, when we choose *ɛ* = 0.015, colonization does not occur until t ≈ 60.

## 3 Discussion and Conclusions

In this work we derive and validate an exact solution for a linear reaction-diffusion PDE on a uniformly growing domain. Our framework is relevant for a general class of uniformly growing domains, 0 < *x* < *L*(*t*), and we present specific results for exponentially-elongating domains, *L*(*t*) = *L*(0)exp(*αt*), with *α* > 0, and linearly-elongating domains, *L*(*t*) = *L*(0)+*bt*, with *b* > 0. While our approach is relevant for a general class of initial conditions, motivated by Landman et al.’s previous work [22], we consider an initial condition relevant to ENS development where we consider *C*(*x*, 0) to be localised near one boundary of the domain. Then, using our exact solution, we explore whether the density profile evolves such that it can overcome the domain growth and colonize the entire length of the domain by reaching the other boundary, within some particular time interval.

It is interesting to note, and discuss, several differences between the solution of the linear reaction-diffusion PDE on a non-growing domain, given by Equation (12), and the solutions of the same PDE on a growing domain, such as Equations (13) and (14). In the usual way, the solution on a non-growing domain (Equation (12)) indicates that after a sufficiently long period of time the exact solution can be approximated by the first few terms in the infinite series since the factor exp 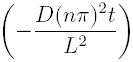 guarantees that further terms in the series decrease exponentially fast with time. On a non-growing domain, this could be used to develop useful approximations to Equation (12), such as

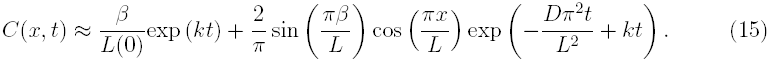

Such approximations are well-known to be accurate after a sufficiently long period of time [25, 26]. One of the key differences between the solutions of Equation (4) on a growing and non-growing domain becomes obvious when we consider whether it is possible to develop a useful approximation of the exact solution in the long-time limit on a growing domain. Since Equation (11) contains the factor exp(—(*nπ*)^2^*T*), it is tempting to think that we may truncate the infinite series after one or two terms to obtain a useful approximation to the exact solution when *T* becomes sufficiently large. This kind of approximation is possible in the non-growing case where, as we previously noted, when *t* → ∞, we have *T* → ∞. However, different behavior occurs in the growing domain solutions. In particular, for the exponentially-growing domain, as *t* → ∞ we have 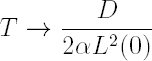. Similarly, in the linearly-growing domain case, as *t* → ∞ we have 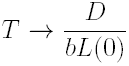. This means that it may not be possible to develop simple approximations for sufficiently large *t*. Indeed, we explored whether it is possible to approximate the exact solutions in Fig. 1 and Fig. 2 using a two-term truncation of Equation (13) and we found that this produced a very poor approximation, even for much larger values of *t* than reported here, such as *t* = 100.

Since we rely on separation of variables and superposition to construct our exact solution, one of the key limitations of our strategy is that the exact solution applies only to a linear reaction-diffusion process. While many reaction-diffusion models are inherently nonlinear, there is a real practical value in the use of linear models, since linear PDE models are often used to approximate the solution of related nonlinear PDE models[27]. For example, Hickson et al. [28] analyses the critical timescale of a nonlinear reaction-diffusion process by arguing that the nonlinear PDE model can be approximated by a linear PDE model. Similarly, Swanson [29] provides key insight into the motion of a moving front of cells by studying an exact solution of a linear PDE model. In this case, Swanson [29] assumes that the linear PDE model can be used to approximate the solution of a nonlinear PDE. Using a similar approach, Witelski [30] studies the motion of wetting fronts in variably saturated porous media, which is governed by a nonlinear PDE, by first analysing the solution a related linear PDE model. These kinds of approximations are invoked in many other situations such as the study of flow in saturated porous media [31, 32], solid-liquid separation processes [33], and certain problems in food manufacturing [34]. Therefore, while our exact solution cannot be applied directly to study the solution of nonlinear PDE models, the basic properties of the linear PDE model can be used to provide insight into reaction-diffusion processes on a growing domain. In addition to this practical value, we believe that our exact solution is also inherently interesting from a mathematical point of view.

There are several ways in which the exact solution strategy presented in this work could be extended. Although we have only considered a single species reaction-diffusion processes with one dependent variable, *C*(*x*,*t*), in principle our solution strategy could also be applied to multispecies reaction-diffusion processes involving several dependent variables, *C*_1_(*x*,*t*), *C*_2_(*x*,*t*), *C*_3_(*x*, *t*),..., that are coupled through a linear reaction network [35,36]. We anticipate that these kinds of multispecies problems could be solved exactly on a uniformly growing domain by first applying a linear transformation which uncouples the reaction network [35]. After this initial uncoupling transformation, our solution strategy could be applied to solve each uncoupled PDE before applying the inverse uncoupling transform to give an exact solution for the coupled multispecies PDE problem on a growing domain. We leave this extension for future consideration.

## Acknowledgments

I acknowledge the Australian Research Council (FT130100148) and am grateful for assistance from Sean McElwain, Scott McCue, Ruth Baker and the referee.

